# A *de novo* chromosome-level genome assembly of *Coregonus sp. “Balchen”*: one representative of the Swiss Alpine whitefish radiation

**DOI:** 10.1101/771600

**Authors:** Rishi De-Kayne, Stefan Zoller, Philine G. D. Feulner

## Abstract

**Background:** Salmonids are of particular interest to evolutionary biologists due to their incredible diversity of life-history strategies and the speed at which many salmonid species have diversified. In Switzerland alone, over 30 species of Alpine whitefish from the subfamily Coregoninae have evolved since the last glacial maximum, with species exhibiting a diverse range of morphological and behavioural phenotypes. This, combined with the whole genome duplication which occurred in the ancestor of all salmonids, makes the Alpine whitefish radiation a particularly interesting system in which to study the genetic basis of adaptation and speciation and the impacts of ploidy changes and subsequent rediploidization on genome evolution. Although well curated genome assemblies exist for many species within the Salmonidae family, genomic resources for the subfamily Coregoninae are lacking.

**Findings:** PacBio sequencing from one wild caught *Coregonus sp. “Balchen”* from Lake Thun was carried out to ∼90x coverage. PacBio reads were assembled independently using three different assemblers, Falcon, Canu and wtdbg2 and subsequently scaffolded with additional Hi-C data. All three assemblies are of high quality based on standard metrics, and when comparing the assemblies to a previously published linkage map and when mapping additional short-read data (∼30x Illumina data) to it.

**Conclusions:** Here, we present the first *de novo* genome assembly for the Salmonid subfamily Coregoninae. Our final wtdbg2 reference sequence was assembled into 40 chromosome-scale scaffolds with a total length of 2.2Gb, an N50 of 51.9Mb and was 93.3% complete for BUSCOs. It comprised of ∼52% TEs and contained 46,397 genes.

## Introduction

Members of the *Coregonus* genus, known as lake whitefish, are distributed throughout freshwater systems across Europe and North America [1–5]. In many lakes across their range, multiple whitefish species have evolved in the last 12,000 years following the melting of glaciers after the last glacial maximum [4–6]. Today a particularly speciose clade of whitefish is found throughout pre-alpine lakes across Switzerland, known as the Alpine whitefish radiation [7]. Over 30 species are thought to make up this radiation which was previously described as the *C. lavaretus spp.* complex, and new studies continue to identify additional cryptic diversity within the radiation using genetic methods [3,5,8,9]. Within Switzerland, independent, monophyletic, radiations of up to six species have evolved rapidly following the last glacial maximum [5,9]. Sympatric whitefish species in these lakes are differentiated in many phenotypic traits including body size, gill-raker number (linked to their feeding ecology) as well as spawning depth and season [4,8,9]. Repeated phenotypic differentiation has evolved independently across different lake systems resulting in allopatric species exhibiting analogous life history strategies e.g. large, shallow spawning, benthic macro-invertebrate eaters *C. sp. “Bodenbalchen” sp. nov., C. sp. “Balchen”* and *C. duplex* are present in lakes Luzern (Reuss system), Thun/Brienz (Aare system) and Walen/Zurich (Limmat system), respectively. Likewise, in the same lakes *C. zugensis, C. albellus*, and *C. heglingus*, small bodied pelagic zooplanktivores with high numbers of gill rakers have also evolved, alongside up to four other sympatric species. This rapid and repeated evolution of multiple whitefish phenotypes and life history strategies has made the Alpine whitefish a particularly interesting system in which to study the genomic basis of adaptation and speciation. The recent use of genomic data gained from reduced representation libraries has demonstrated the power of genomic approaches for species designation amongst closely related sympatric species [10]. Further, it was demonstrated that genetic differentiation across the genome is widespread when comparing sympatric species from contrasting habitats [10]. However, the low density and uncertainty of positioning of markers along the genome currently still limits a true genome-wide view of adaptation and speciation within these radiations.

The Salmonidae are a focal family in which to study genome evolution, specifically the rediploidization process following whole genome duplication. As part of the Salmonidae family, Coregonids share a common ancestor with the Salmoninae and Thymallinae. Before these sub-families split from one another, the whole lineage experienced a whole genome duplication 80-100 Mya [11–13]. Recent studies have determined that different Salmonid genomes are uniquely shaped by rediploidization following this whole genome duplication, referred to as the salmonid-specific fourth vertebrate whole-genome duplication, Ss4R [14]. It has been shown that whilst many regions of Salmonid genomes rediploidized prior to the diversification of the three sub-families, and thus are shared across the family, each lineage also has unique patterns of rediploidization for some genomic regions leading to substantial variation in genome structure between lineages [14]. To fully understand the impact of whole genome duplication and subsequent rediploidization on genome structure in the Salmonidae, high quality genome assemblies for all major lineages are needed.

Although many salmonid species now have suitably well assembled and curated reference genomes, including Atlantic Salmon (*Salmo salar*; [13]), Rainbow Trout (*Oncorhynchus mykiss*; [15]), Chinook Salmon (*Oncorhynchus tshawytscha*; [16]), Coho Salmon (*Oncorhynchus kisutch*; NCBI BioProject: PRJNA352719), Arctic Charr (*Salvelinus alpinus*; NCBI BioProject: PRJNA348349; [17]), and Grayling (*Thymallus thymallus*; [18,19]), genomic resources for the Coregoninae subfamily are largely limited. To date, the best curated genomic resources for the Coregoninae are two next-generation sequencing linkage maps, one for the North American system [20] and one for the Alpine whitefish radiation [21].

To enable future studies to investigate both the genetic basis of adaptation and speciation in Alpine whitefish and genome evolution following whole genome duplication across the Salmonidae family we have assembled the first whitefish reference genome. Assembling > 90x PacBio data from one female *Coregonus sp. “Balchen”* with three of the most commonly used assemblers resulted in three high quality assemblies, each with >90% complete BUSCOs and 40 chromosome-scale scaffolds. Out of these three assemblies we judge the assembly produced by wtdbg2 as the best. This new draft whitefish genome is 2.2 Gb and comprises 7,855 scaffolds, including 40 chromosome level scaffolds containing 94.02% of nucleotides, has an N50 of 51.9 Mb and contains 93.3% complete BUSCOs. Annotation of the assembly identified 46397 genes in total and showed that transposable elements make up 52% of the *C. sp. “Balchen”* genome.

## Data description

### Sample preparation and sequencing

DNA was extracted in multiple batches from heart and somatic muscle tissue of one wild caught (outbred) female *C. sp. “Balchen”* from Lake Thun (in December 2016) using the MagAttract HMW DNA Kit (Qiagen). From this high molecular weight DNA, 8mg was used to prepare three libraries for sequencing on the single-molecule, real-time sequencing (SMRT) platform from Pacific Biosciences using 60 SMRT cells to generate 383 gigabytes (240 Gb) of data (Next Generation Sequencing Platform, University of Bern). Illumina sequencing of paired end reads (of 150 bp) was also carried out using the Hi-Seq platform (Next Generation Sequencing Platform, University of Bern) which generated 87 Gb of data to be used for polishing.

### Estimation of genome size

Reports of *Coregonus* genome sizes vary widely with estimates ranging from 2.4 Gb [22] to 3.5 Gb [23]. To better estimate genome size for our focal species *Coregonus sp. “Balchen”*, we used Jellyfish v.1.1.11 [24] to produce frequency distributions of 17-, 21-, 25 and 30-mers for all Hi-Seq reads. GenomeScope v.1 [25] was then used to estimate genome size from these histograms. The best model (k=25), as determined using the ’Model Fit’ output from GenomeScope estimated *C. sp. “Balchen”* to have a genome size of 2.63 Gb (Figure 1). Based on this estimate of genome size our PacBio sequencing equates to ∼91x coverage (and our Illumina Hi-Seq ∼33x).

**Figure 1.**
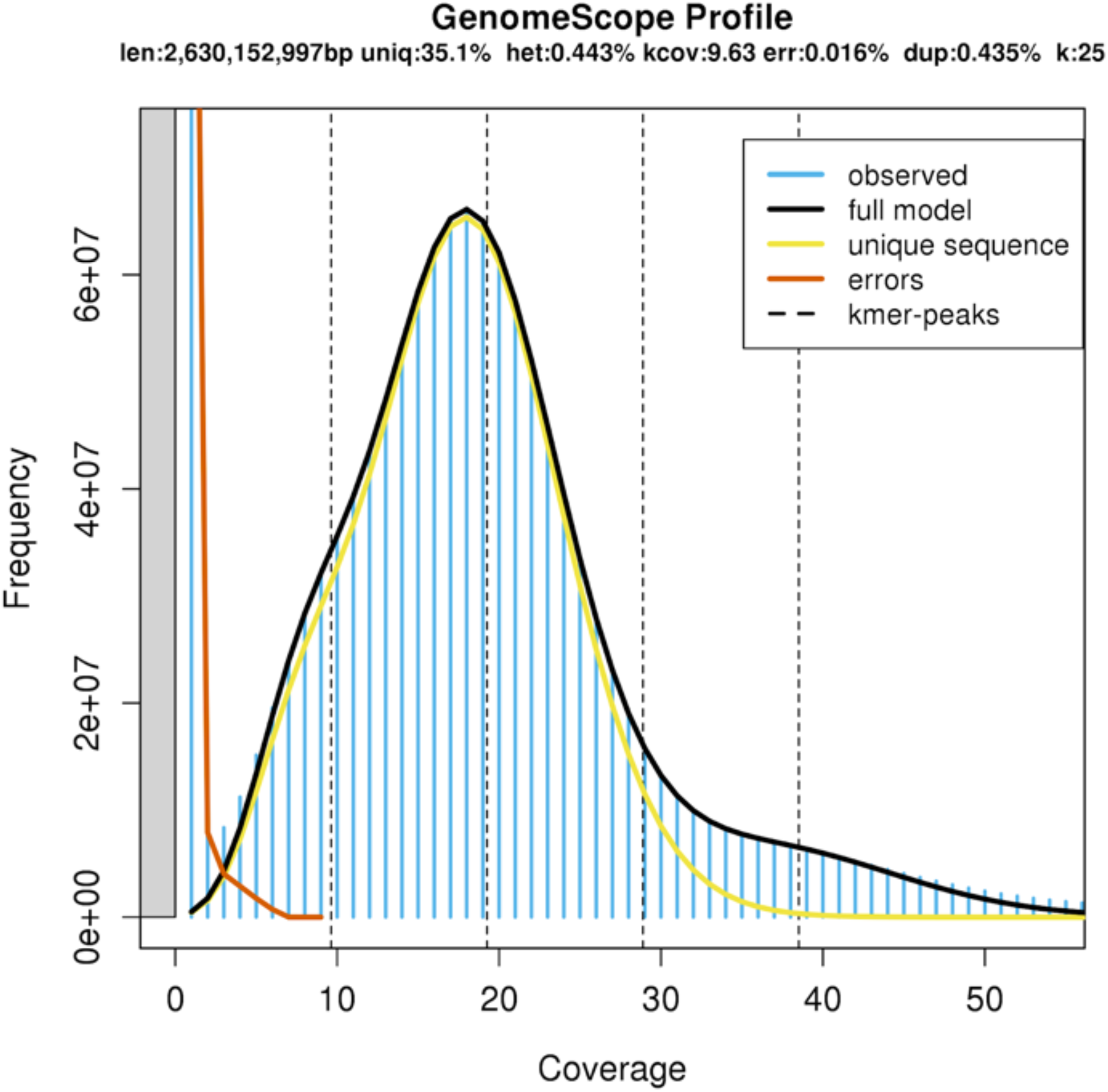
GenomeScope profile established based on short read data, which estimates the genome size of *Coregonus sp. “Balchen”* to be 2.6 Mb.

### Genome assembly and polishing

Raw PacBio data was assembled independently using three different assemblers (Figure 2), Falcon/Falcon Unzip [26], Canu [27], and wtdbg2 [28], which have each demonstrated their ability to produce highly contiguous assemblies. In all three cases assembly was carried out using only PacBio data. Read polishing was carried out using both the original raw PacBio reads (Arrow; [29]) and low-error rate, short-read, Illumina data (Pilon; [30]). After each round of polishing BUSCO v.3.0.2 [31] was run against the core gene set from ray-finned fishes (actinopterygii_odb9) to evaluate the assembly. If the number of complete BUSCOs did not improve after running the Arrow algorithm then only Pilon was used. All assembly parameters can be found at https://github.com/RishiDeKayne/WhitefishReferenceGenome.

**Figure 2.**
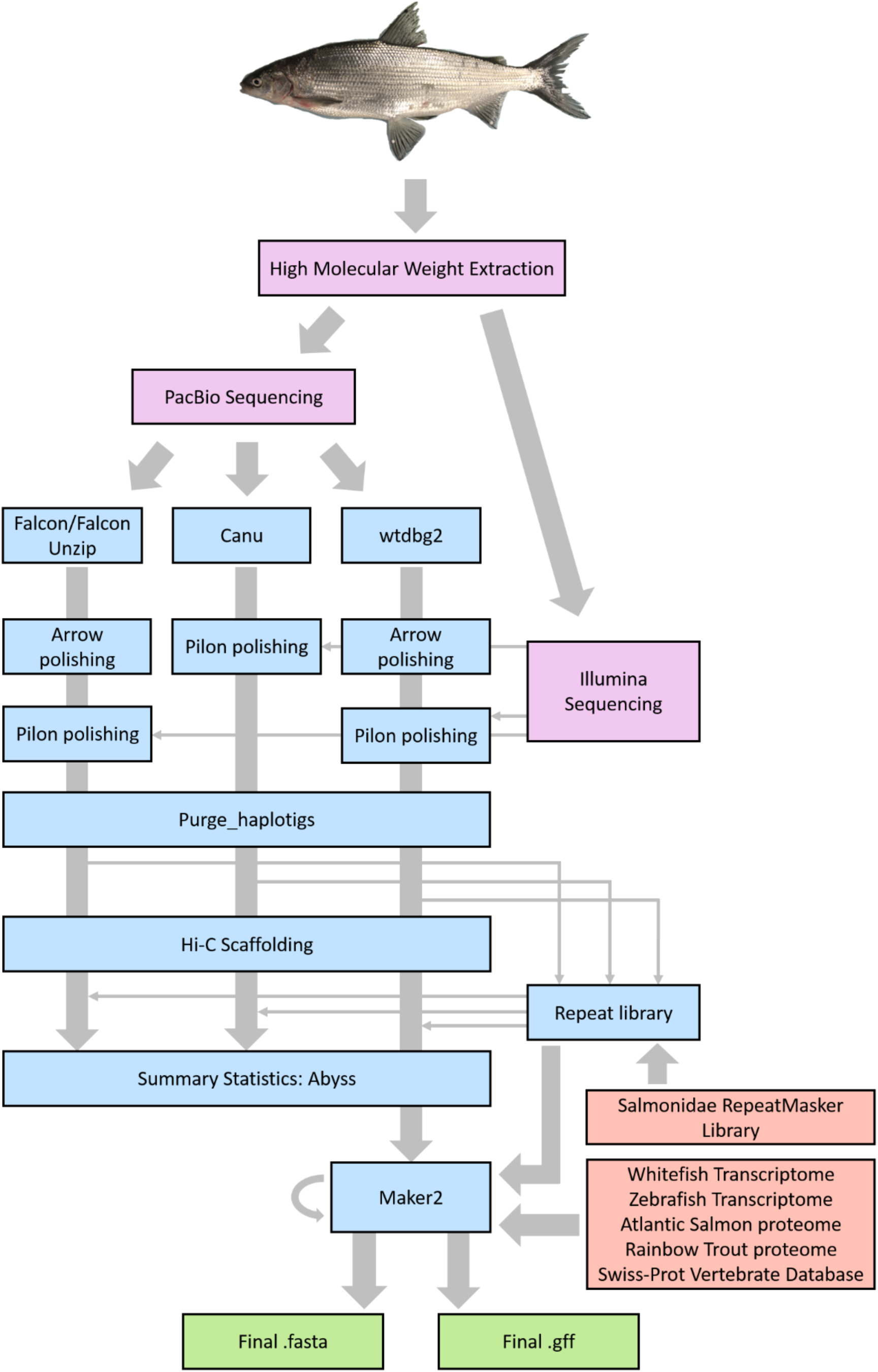
Workflow outlining the different steps and tools used to assemble the whitefish genome (coloured in blue). New input produced for this study is coloured in purple and previously published resources used for repeat masking and annotation in orange. Final outputs are shown in green.

#### Falcon/Falcon Unzip

Genome assembly was carried out by DNAnexus (Mountain View, CA) utilizing the DAMASKER suite [32] and the FALCON (v1.9.1) pipeline (Pacific Biosciences; [26]). First the REPmask and TANmask modules of the DAMASKER suite were used on the raw PacBio reads and the resulting output was used as input for the FALCON 1.9.1 pipeline. For the first two steps of the Falcon pipeline, error-correction and read overlap identification, a length cut-off of 5000 bp was used. These aligned reads were then assembled into 52,448 primary contigs containing 2.78 Gb with an N50 contig length of 204 Kb. This assembly was then phased and polished using Falcon-Unzip [26] and Pacific Biosciences’ Arrow algorithm [29]. The final step involved polishing these contigs using ∼33x Illumina reads in the Pilon program v.1.22 [30]. This resulted in primary contigs, thought to represent the haploid whitefish genome, and haplotig contigs, thought to represent alternative alleles at heterozygous sites in our subject fish. The primary contigs assembly was made up of 19,553 contigs covering 2.41 Gb and with an N50 of 280 Kb (Table 1). For downstream processing of the Falcon assembly this primary contigs assembly was combined with the reads identified as haplotigs by Falcon Unzip. This allowed us to find misidentified primary contigs (which may rather represent haplotigs or mitochondrial DNA) as well as misidentified haplotigs (which could in fact be low coverage contigs or repetitive/duplicated regions). This merged assembly was 4.11 Gb long, contained 60,605 contigs and included 89.5% of complete BUSCOs (Table 1; Table S1).

**Table 1.** Summary statistics at the contig (pre-haplotig purging) and scaffold stage (each scaffolded assembly contains 40 chromosome-scale scaffolds) for the Falcon, Canu and wtdbg2 assemblies.

|  |  | **Falcon (primary contigs)** | **Canu** | **wtdbg2** |
| --- | --- | --- | --- | --- |
| Contig statistics |  |  |  |  |
|  | Number of contigs | 60605 (19553) | 52023 | 28224 |
|  | Contig N50 (bp) | 136418 (279657) | 130955 | 424474 |
|  | Longest contig (bp) | 6516619 (6516619) | 5278180 | 5201837 |
|  | total contig length (Gb) | 4.11 (2.41) | 3.28 | 2.38 |
| Scaffold statistics |  |  |  |  |
|  | Number of scaffolds | 3745 | 3553 | 7855 |
|  | Scaffold N50 (bp) | 62840000 | 59340000 | 51930000 |
|  | Longest scaffold (bp) | 111300000 | 104000000 | 93420000 |
|  | total scaffold length (Gb) | 2.47 | 2.46 | 2.20 |
| All scaffold BUSCOs/<br>chromosome-scale scaffold BUSCOs |  |  |  |  |
|  | Complete | 4209 (91.8%)/4195 (91.5%) | 4299 (93.7%)/4297 (93.7%) | 4274 (93.3%)/4263 (93%) |
|  | Single | 2713 (59.2%)/2732 (59.6%) | 2578 (56.2%)/2583 (56.3%) | 2551 (55.7%)/2551 (55.7%) |
|  | Duplicated | 1496 (32.6%)/1463 (31.9%) | 1721 (37.5%)/1714 (37.4%) | 1723 (37.6%)/1712 (37.3%) |
|  | Fragmented | 89 (1.9%)/77 (1.7%) | 78 (1.7%)/78 (1.7%) | 95 (2.1%)/83 (1.8%) |
|  | Missing | 286 (6.3%)/312 (6.8%) | 207 (4.6%)/209 (4.6%) | 215 (4.6%)/238 (5.2%) |

#### Canu

Assembly following the Canu (v1.6) pipeline [27] was carried out using the same raw PacBio data. Canu assembly includes three main steps, error correction followed by read trimming and finally, assembly. Canu read correction was carried out using the default settings regarding minimum read length (1000 bp) and minimum read overlap (500 bp) whilst specifying a genome size of 4 Gb. The same parameters were used for the trimming step however, for the assembly step minimum read length was extended to 1200 bp and minimum read overlap to 600 bp. Similar to the Falcon/Falcon Unzip assembly, the final step involved polishing contigs using ∼33x Illumina reads in the Pilon program v.1.22 (one round; [30]). The resulting Canu assembly was substantially larger than the primary reads from the Falcon/Falcon Unzip assembly covering 3.28 Gb across 52,023 contigs, with an N50 of 131Kb and including 88.7% BUSCOs (Table 1; Table S1).

#### wtdbg2

Finally, an assembly was carried out with the least computationally intensive of the three programs -wtdbg2 [28]. wtdbg2 involves two steps, the first of which assembles long reads and the second derives a consensus sequence. For long read assembly kmer psize was set to 21 (-p 21), and 1/3 kmers were subsampled (using -S 3), the maximum length variation of two aligned fragments was set to 0.05 (-s 0.05) and the minimum length of alignment was 5000 bp (-L 5000). After the consensus was derived one round of polishing was carried out using Arrow followed by two rounds of polishing with Pilon v.1.22 [30]. This wtdbg2 assembly was the shortest of the three assemblies with a total length of 2.38 Gb and also had the fewest contigs (28,224 Table 1). However, it had the highest N50 of 424 Kb and contained the highest percentage of complete BUSCOs, 93.4% (Table 1; Table S1).

### Haplotig purging

Following each assembly, we used purge_haplotigs [33] to identify contigs that were more likely to represent alternative alleles (from heterozygous regions of the genome) or mitochondrial DNA rather than the haploid nuclear genome. In each case, all raw PacBio data was mapped against the assembly and a read-depth histogram was produced. A low, mid, and high value of coverage was then selected from this histogram to flag suspect haplotigs and regions with exceptionally high coverage (all thresholds and histograms can be found in Table S1). Suspect haplotigs were then mapped to the rest of the assembly to identify their allelic partner before the contigs with good matches were reassigned as haplotigs. After haplotig purging the differences between the three assemblies was reduced dramatically with the range of contigs now from 16,440 to 22,627 (for wtdbg2 and Canu, respectively; Table S1). The N50 of all three assemblies also increased, substantially in the Falcon and Canu assemblies from 136 Kb and 131 Kb to 281 Kb and 258 Kb each. The N50 of the wtdbg2 assembly also increased from 424 Kb to 491 Kb (Table S1). To assess the gene-level completeness of each assembly after running purge_haplotigs each assembly was again compared against the core gene set from ray-finned fishes (actinopterygii_odb9) using BUSCO v.3.0.2 [31]. The number of complete BUSCOs went up in both the Falcon and Canu assemblies (by 1.3% and 4.4%) but dropped slightly (by 0.3%) in the wtdbg2 assembly (Table S1). The high completeness percentage of BUSCOs for each of our assemblies prior to scaffolding suggests that we have succeeded in capturing a large proportion of the whitefish genome sequence during the assembly process.

### Genome scaffolding

Hi-C scaffolding of the purged assemblies, including tissue processing, library preparation, and sequencing, was carried out by Phase Genomics (Seattle, WA).

Chromatin conformation capture data was generated using a Phase Genomics (Seattle, WA) Proximo Hi-C Animal Kit, which is a commercially available version of the Hi-C protocol [34]. Following the manufacturer’s instructions for the kit, intact cells from the same whitefish female were crosslinked using a formaldehyde solution, digested using the Sau3AI restriction enzyme, and proximity ligated with biotinylated nucleotides to create chimeric molecules composed of fragments from different regions of the genome that were physically proximal *in vivo*, but not necessarily genomically proximal. Continuing with the manufacturer’s protocol, molecules were pulled down with streptavidin beads and processed into an Illumina-compatible sequencing library. Sequencing was performed on an Illumina HiSeq 4000, generating a total of 249,544,461 100 bp read pairs.

Following the manufacturer’s recommendations [35], reads were aligned independently to each of the three draft assemblies (Canu, Falcon and wtdbg2). Briefly, reads were aligned using BWA-MEM [35] with the -5SP and -t 8 options specified, and all other options default. SAMBLASTER [36] was used to flag PCR duplicates, which were later excluded from analysis. Alignments were then filtered with samtools [37] using the -F 2304 filtering flag to remove non-primary and secondary alignments and matlock [38] using default options. Putative misjoined contigs were broken using Juicebox [39] based on the Hi-C alignments; 11 breaks were introduced into the Canu assembly, 42 breaks into the Falcon assembly, and 11 breaks into the wtdbg2 assembly.

Phase Genomics’ Proximo Hi-C genome scaffolding platform was used to create chromosome-scale scaffolds from the draft assembly in a method similar to that described by Bickhart et al. [40]. As in the LACHESIS method [41], this process computes a contact frequency matrix from the aligned Hi-C read pairs, normalized by the number of Sau3AI restriction sites (GATC) on each contig, and constructs scaffolds in such a way as to optimize expected contact frequency. For each of the four assemblies, approximately 100,000 separate Proximo runs were performed to optimize the number of scaffolds and the scaffold construction in order to make the scaffolds as concordant with the observed Hi-C data as possible. This process resulted in a set of 40 scaffolds for each of the three assemblies, containing 2.38 Gb (96.5% of all sequence; Falcon), 2.41 Gb (97.8% of all sequence; Canu) and 2.07 Gb (94.0% of all sequence; wtdbg2). Finally, each set of scaffolds was polished an additional time using Juicebox [39].

The differential log-likelihood of each set of scaffolds was calculated and examined in the same manner demonstrated by LACHESIS. A threshold of 100 was used to identify contigs scaffolded in a position and orientation in which the log-likelihood (base e) of the chosen orientation was more than 100 times greater than the alternative, a method shown by Burton et. al [41] to be effective for identifying contigs that are well ordered and oriented in their region of a scaffold. The length of sequence meeting this criterion was 1.08 Gb (45.41%) for the Falcon scaffolds, 1.21 Gb (50.1%) for the Canu scaffolds, and 1.74 Gb (84.1%) for the wtdbg2 scaffolds. These results are in agreement with the patterns observable in the final scaffold heatmaps for each, in which the patterns observable for the wtdbg2 scaffolds are more in alignment with a priori expectations about Hi-C linkage density patterns (Figure 3; [42]), yielding qualitative confirmation of the quantitative scaffold quality assessment. Following scaffolding each of the assemblies, BUSCO v.3.0.2 [31] was run again on each complete assembly as well as the 40 largest, chromosome-scale scaffolds. The percentage of complete BUSCOs went up for each assembly with the Falcon, Canu, and wtdbg2 assemblies now having 91.8%, 93.7% and 93.3% (Table 1). When considering only chromosome-level scaffolds the Canu assembly retained the highest complete percentage of BUSCOs at 93.7% with the Falcon and wtdbg2 assemblies dropping only slightly to 91.5% and 93.0% each. Based on the high proportion of the contigs confidently scaffolded and the high number of complete BUSCOs, the Hi-C scaffolded wtdbg2 assembly was selected as the best of the three assemblies and was uploaded to the European Nucleotide Archive (Accession: ERZ1030224). Falcon and Canu assemblies are available on request.

**Figure 3.**
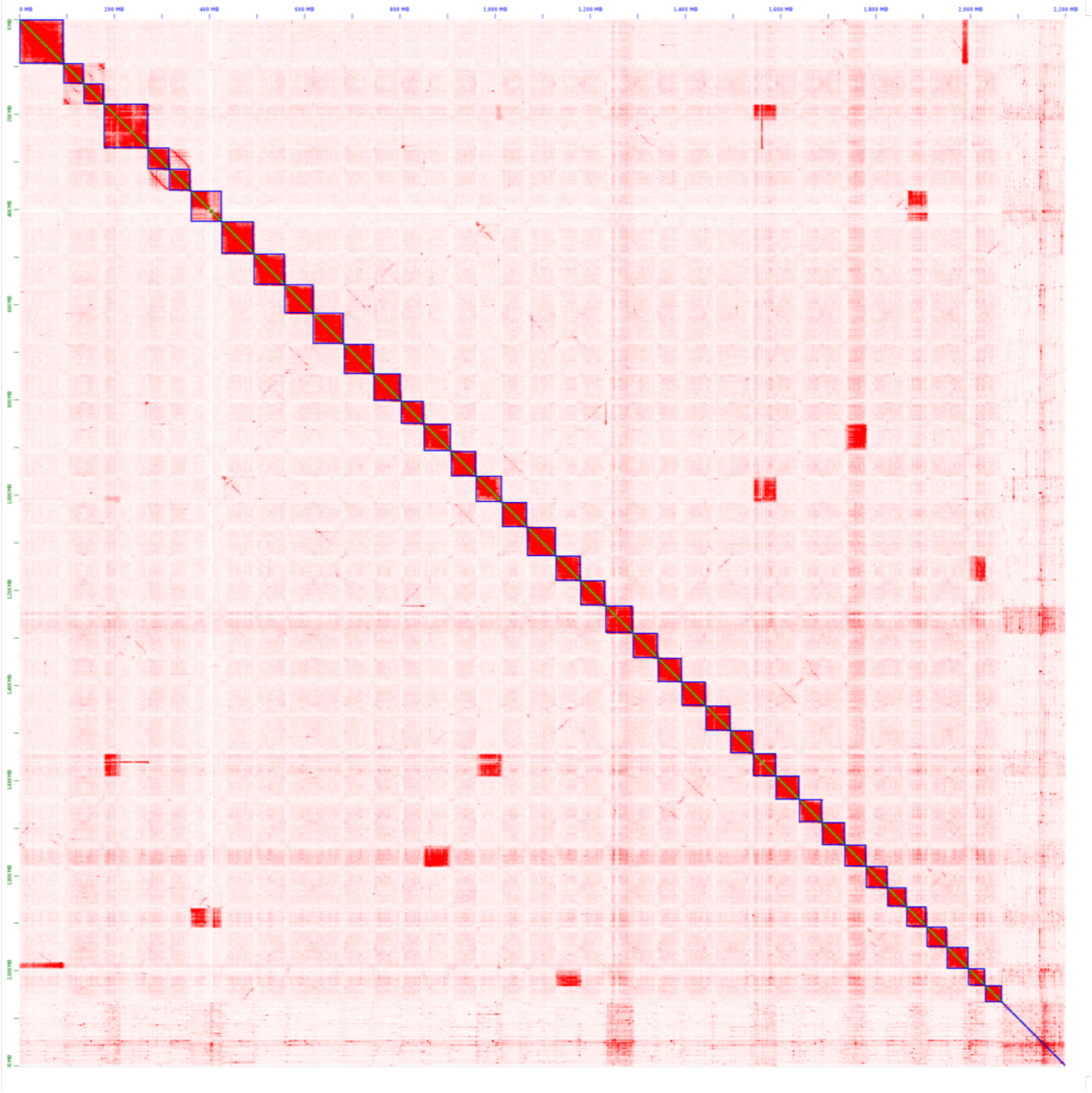
*Coregonus sp. “Balchen”* contig contact map from Hi-C scaffolding of the wtdbg2 assembly. The intensity of red represents the relative contact density between contigs (green along the diagonal). The highest contact density is found within scaffolds, which are outlined in blue.

### Validation of chromosome level assemblies

#### Illumina short read mapping

To assess the qualitative differences between the three scaffolded assemblies we used two independent data sets, the Illumina short reads and a previously published *Coregonus sp. “Albock”* linkage map (see next section). Mapping the Illumina data helped to assess the composition of each of the chromosome-scale scaffolds. The Illumina data, collected from the same individual from which the genome was sequenced, was mapped back to each of the reference assemblies. In this way we assessed the consistency of coverage across the assembly and identify potentially duplicated regions of the whitefish genome which may have been collapsed into one sequence during the assembly process. Illumina reads were mapped to each assembly using BWA-MEM ([35]; with default parameters). A summary of this mapping was produced using samtools ([37]; samtools flagstat). Coverage was then calculated in 30 Kb windows using bedtools [43] and a custom perl script cov.per.window.pl. Coverage statistics were then calculated in R [44].

Summaries of the mapping of Illumina reads to each of the assemblies can be seen in Table S2. The wtdbg2 assembly had the highest number of mappings over MAPQ 30 (71.6%) with the Canu and Falcon assemblies having slightly lower proportions of high-quality mappings (69.5% and 60.7% each). When considering the proportion of read mates mapped to a different chromosome, however, the Canu assembly looks the best of the three with only 1.4% of mates mapped to a different chromosome with a MAPQ > 5, compared to 2% for Falcon and 2.4% for wtdbg2. As a result of our coverage analysis the highest mean coverage (in 30 Kb windows) was 17.3x observed in the wtdbg2 assembly. The Falcon and Canu assemblies had lower mean coverages of 15.0x and 15.9x, respectively. Plots of coverage across the 40 chromosome-scale scaffolds are shown in Figure 4. Based on these coverage plots we identified regions which are likely to represent collapsed duplicated regions, spread across each genome assembly. As expected in these regions, the coverage was approximately double that of the rest of the assembly. In other salmonid genome assemblies, which have successfully resolved each copy of a duplicated region, these duplicated regions typically span whole chromosome arms or even chromosomes. Similarly, in our assemblies we identify collapsed blocks which encompass whole chromosome-level scaffolds or large proportions of chromosome-level scaffolds. In the wtdbg2 assembly some scaffolds (scaffolds 3, 6, 16 and 36) may represent collapsed regions which span only chromosome arms, and in other scaffolds (scaffolds 21, 27, 31, 34, and 37) the whole chromosome looks collapsed (a BED file containing the locations of these collapsed duplicates is included as Supplementary file S3).

**Figure 4.**
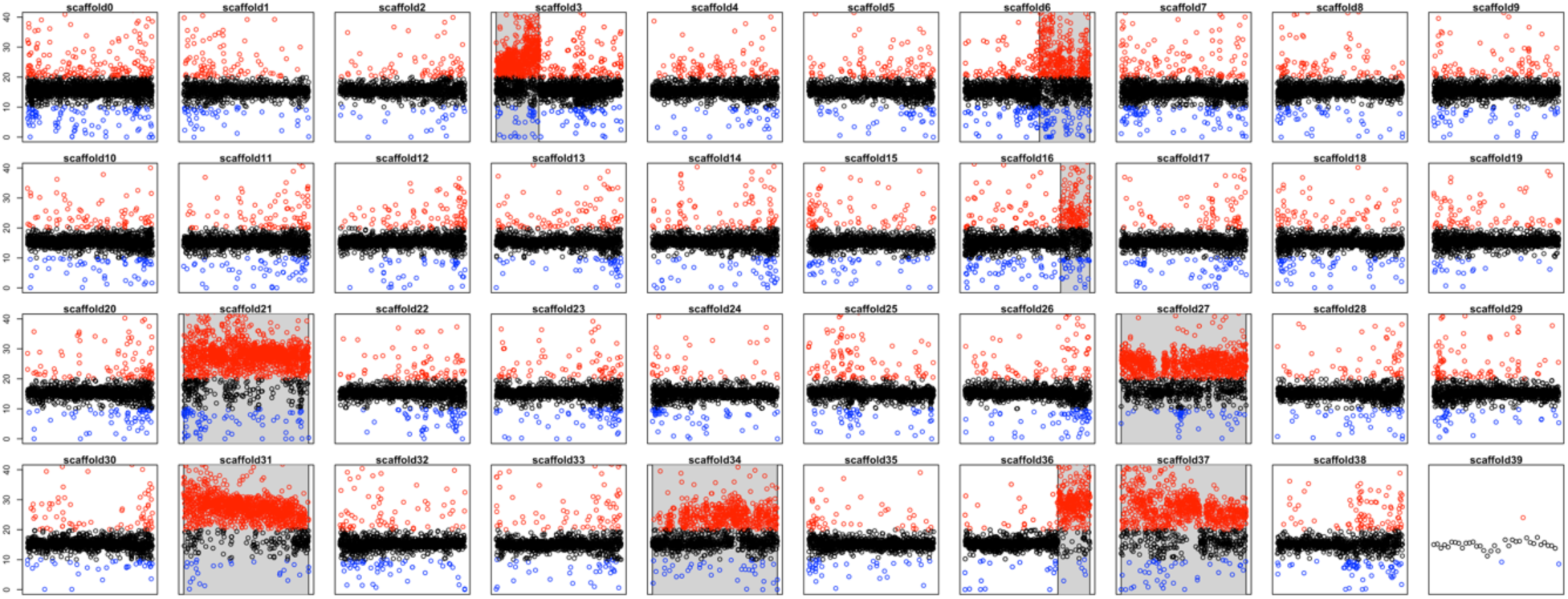
For each of the chromosome-scale scaffolds coverage of Illumina data mapped to the wtdbg2 assembly is plotted in 30 Kb windows. Most windows show an average coverage of around 17x (black points). Windows with coverage >20, and <10 are coloured in red and blue respectively. Putative collapsed duplicate regions are highlighted in grey.

#### Linkage map synteny

In addition, we were able to assess the reliability of scaffolding by investigating the synteny between the 40 chromosome level scaffolds from each assembly and the *C. sp. “Albock”* linkage groups [21]. RAD loci (90 bp containing a marker) with a known position in the linkage map were mapped to the 40 chromosome-scale scaffolds from each assembly using BWA-MEM ([35]; with default parameters).

Out of the 5395 markers only high-quality mappings (MAPQ > 30) were retained, resulting in a mapping position of 3648, 4494, and 4744 markers in the sequence of the Falcon, Canu, and wtdbg2 assemblies respectively (Table S2). For all three assemblies concordance between sequence and recombination position across the majority of markers was very high, suggesting a high synteny between linkage groups and chromosome-scale scaffolds. In the wtdbg2 assembly 95% of markers (4489/4744) showed strong synteny between one linkage group and one chromosome-level scaffold (38/40 linkage groups; Figure 5). Only two scaffolds, (scaffolds 37 and 39) could not be matched to any linkage group. We also identified a series of substantial deviations from the broader pattern, where a number of markers from a linkage group also mapped to a second, alternative scaffold. This was the case for markers from Calb01 – scaffold 36, Calb02 – scaffold 31, Calb08 – scaffold 37, Calb13 – scaffold 34, Calb16 – scaffold5, Calb20 – scaffold 27, Calb34 – scaffold 6, and Calb36 – scaffold 3. Strikingly the mapping location of seven of these deviations (scaffolds 3, 6, 27, 31, 34, 36, and 37) also represent seven of the nine scaffolds we identified as collapsed duplicates showing inflated coverage (shown in grey in Figures 4 and 5). Although scaffold 16 looked like a collapsed duplicate, no significant deviations of markers mapped to this scaffold. Additionally, despite having an unusual mapping pattern with markers from Calb16 in addition to those from Calb33 scaffold 5 showed a consistent coverage of Illumina reads mapping. Out of all deviating markers, 83% (212/255) mapped to regions identified as collapsed duplicates. No markers from the linkage map were successfully mapped to scaffold 39, the smallest of the ’chromosome-level’ scaffolds, at only 1.1 Mb long. Additionally, in two cases markers from two linkage groups predominantly mapped to one wtdbg2 scaffold. Markers from both Calb35 and Calb40 mapped to scaffold 30 and from Calb38 and Calb39 to scaffold 21.

**Figure 5.**
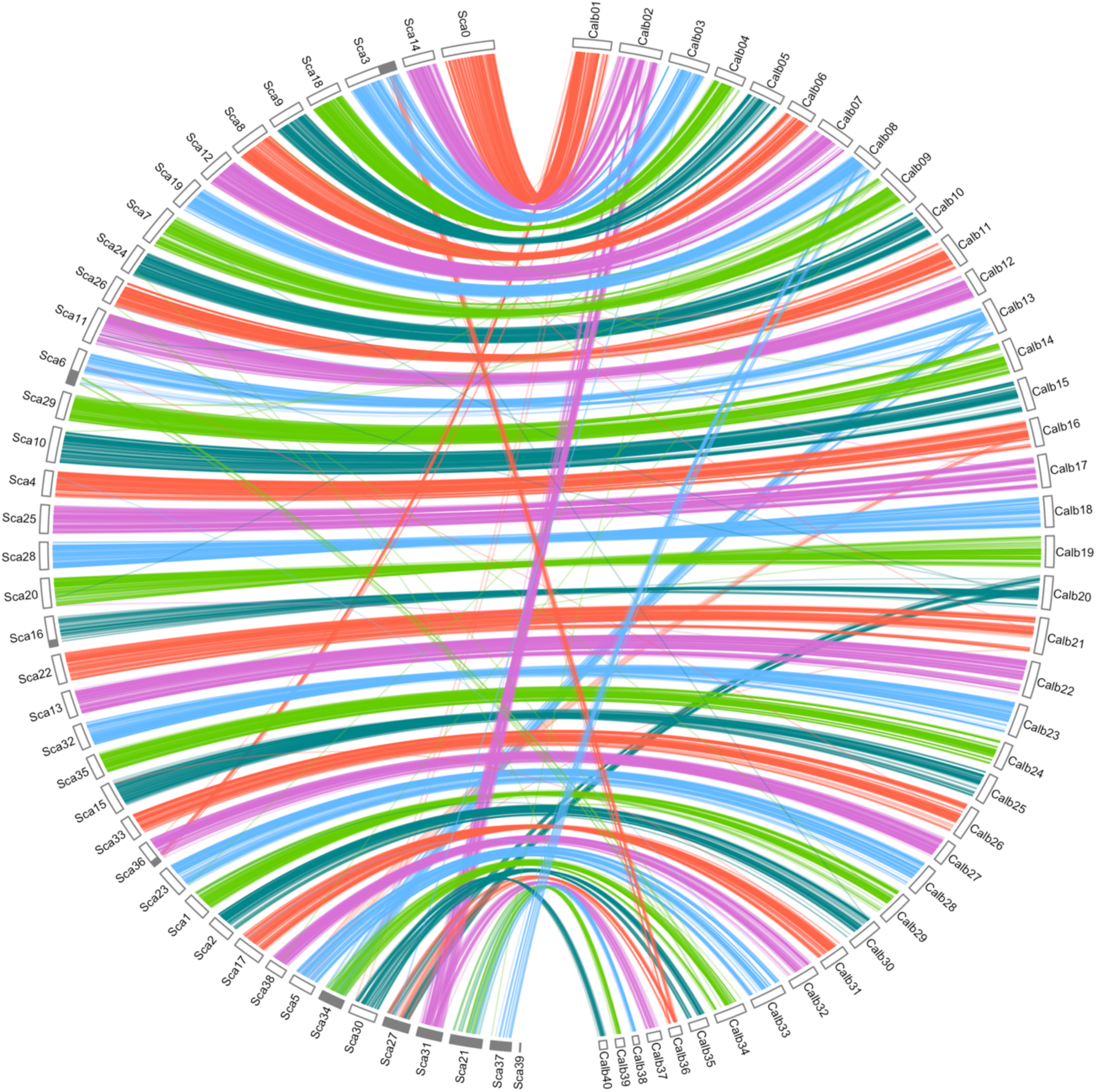
Circos plot comparing the structure of the *C. sp. “Albock”* linkage map (right; [21]) and the 40 chromosome-scale scaffolds of the wtdbg2 *C. sp. “Balchen”* assembly (left). Lines indicate mapping locations of RAD loci from the linkage map in the genome assembly. Most mappings suggest a good match between linkage map and genome assembly (high synteny between linkage groups and chromosomes) and only few lines map discordantly. Genome assembly regions which represent collapsed duplicate regions are identified in grey around the left perimeter.

### Repeat masking

To characterize the repeat landscape of the whitefish genome we first produced a repeat library using RepeatModeler v1.0.11 [45] for each of the haplotig-purged assemblies. These libraries were then combined with the Salmonidae Repbase library (queryRepeatDatabase.pl -species Salmonidae) to produce one concatenated reference library. Each of the scaffolded assemblies were then repeat masked using this concatenated library with RepeatMasker v4.0.7 [46]. An interspersed repeat landscape was then produced for the wtdbg2 assembly using the RepeatMasker scripts calcDivergenceFromAlign.pl and createRepeatLandscape.pl.

Around 52% of each assembly was masked with the most abundant repetitive elements being DNA elements followed by LINEs and then unclassified repeats (Table S1). DNA elements alone made up nearly a quarter of each assembly (24.65% of Falcon, 23.79% of Canu, and 24.41% of the wtdbg2 assembly). The relative abundances of each of these types of repetitive element are similar to those reported in other salmonid assemblies including that of Atlantic Salmon [13] and Chinook Salmon [16]. The resulting landscape (Figure 6) identified the Class II TE superfamily Tc1-*mariner* as the most abundant in the *C. sp. “Balchen”* genome, making up 18% of the interspersed repeats. The most abundant Class I TEs were LINE-2 elements, however, these only made up 4.2% of the interspersed repeats. The relatively high abundance of Class II TE superfamily Tc1-mariner and LINE-2 elements amongst the youngest elements suggest that these families are still expanding and potentially diversifying in the whitefish genome. Conversely, the lack of new LTR elements suggests that their abundance and diversity peaked in the past and they are no longer diversifying in the genome.

**Figure 6.**
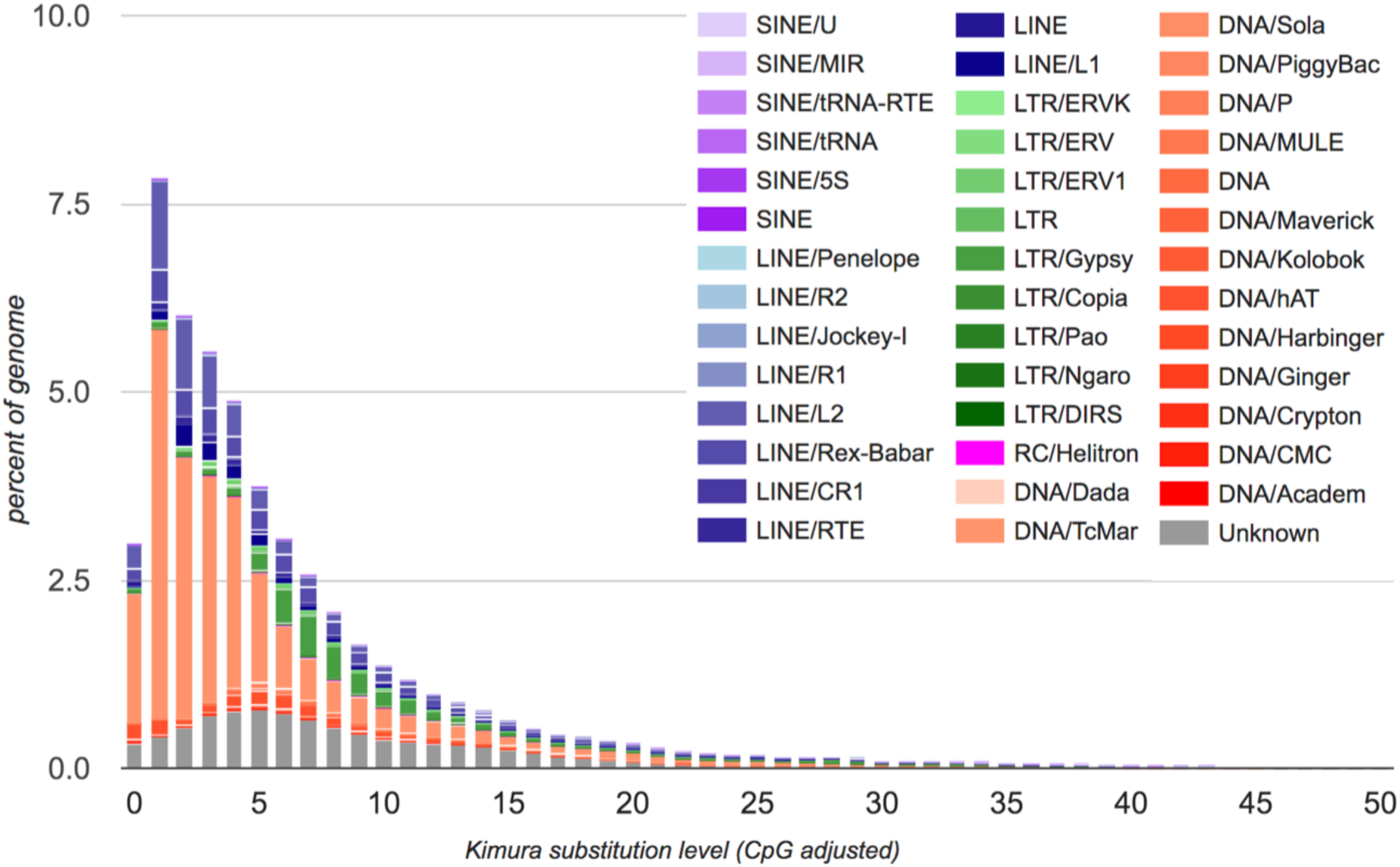
*Coregonus sp. “Balchen”* transposable element divergence landscape. Transposable elements within the whitefish genome have been characterised (different classes represented by distinct colours). The plot shows the relative abundance of each class and their relative age (molecular clock estimate). Note the ongoing DNA element diversification within the whitefish genome, particularly in DNA elements and LINEs.

### Genome annotation

Annotation of the wtdbg2 assembled genome was carried out using a three-pass iterative approach with MAKER2 v2.31 [47]. First, an initial gene model was made using our repeat library (described above), protein evidence from *Salmo salar* (UPID: UP000087266) and *Oncorhynchus mykiss* (UPID: UP000193380) proteomes from Uniprot and the Swissprot vertebrate database (uniprot_sprot_vertebrates), a recently published whitefish transcriptome [48] and alternative transcriptome evidence from a *Danio rerio* transcriptome (TSA: GDQQ01000001:GDQQ01083602). Next, this gene model was used to produce HMMs with SNAP [49] and Augustus v3.2.1 [50]. A second pass of MAKER2 was then carried out using these *ab initio* gene prediction models and the models were optimized before a final third MAKER2 pass was carried out. These gene predictions resulted in 46,397 protein-coding genes (Table 2). Finally, functional annotation was carried out using Pannzer2 [51] resulting in 34,697 genes with an associated gene ontology term.

**Table 2.** Genome annotation summary statistics for final wtdbg2 assembly following three-pass MAKER2 annotation.

|  |  |  |
| --- | --- | --- |
| Genes | Number | 46397 |
|  | Mean length (bp) | 11545.9 |
|  | Median length (bp) | 206 |
|  | Min/max (bp) | 77/187074 |
|  | Gene frequency<br>(genes/Mb) | 21.10 |
| Exons | Number | 365648 |
|  | Mean length (bp) | 196.793 |
|  | Median length (bp) | 164 |
|  | Min/max (bp) | 2/17274 |

In conclusion we have assembled a chromosome-level reference genome for the Coregoninae subfamily. To obtain the most complete whitefish assembly we tested three of the most widely used assemblers. We compared summary statistics, BUSCOs, evenness of coverage from additional short read data, and synteny to a published linkage map amongst the assemblies. We determined that PacBio sequence data acquired for a female *Coregonus sp. “Balchen”* was best assembled by wtdbg2. We were further able to scaffold the resulting contigs using Hi-C to obtain an assembly resulting in 40 chromosome-scale scaffolds and 7,815 contigs with an N50 of 51.9 Mb. In total the assembly contained 46,397 genes and was rich in TEs with a diversity typical for salmonids. This assembly will facilitate future research into both the genetic basis of repeated parallel adaptation and speciation we observe in Alpine whitefish as well as the consequences of whole genome duplication across the whole Salmonidae family.

## Acknowledgements

Thanks to Benjamin Gugger and team from Lake Thun whitefish hatchery for providing us with the *C. sp. “Balchen”* used for genome assembly. Also, thanks to Daniel Jeffries for many helpful discussions regarding our genome assembly pipeline, Oliver Selz for his taxonomic expertise and Cameron Hudson for his suggestions to improve the clarity of the manuscript. Thanks to the Triple A Winter School 2017 (ETH Zurich) for providing invaluable core skills for the project. Data produced and analysed in this paper were generated in collaboration with the Next Generation Sequencing Platform, University of Bern and the Genetic Diversity Centre (GDC), ETH Zurich. This project is funded by the Swiss National Science Foundation (SNSF project 31003A_163446/1 awarded to PGDF).

---

**Table S1.**
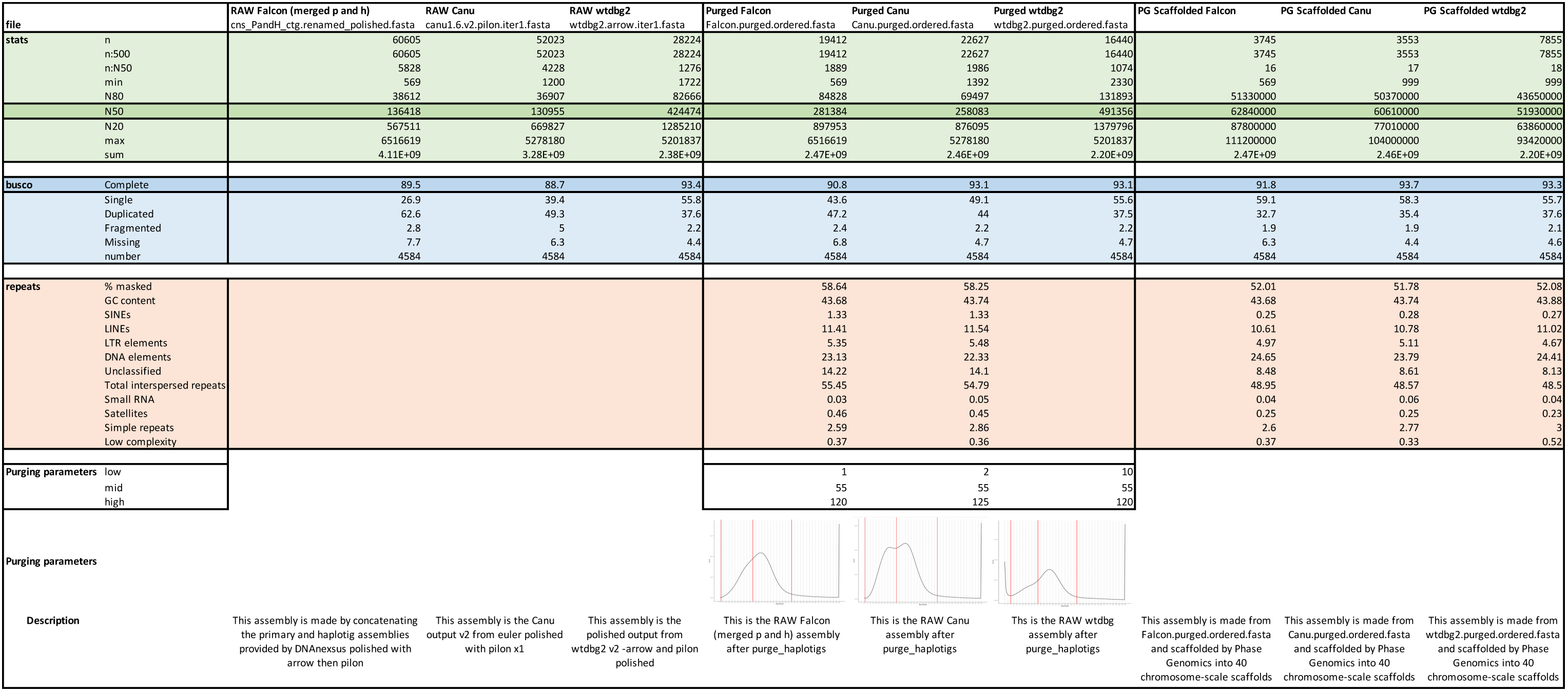
Summary statistics at each stage of the assembly process

**Table S2.**
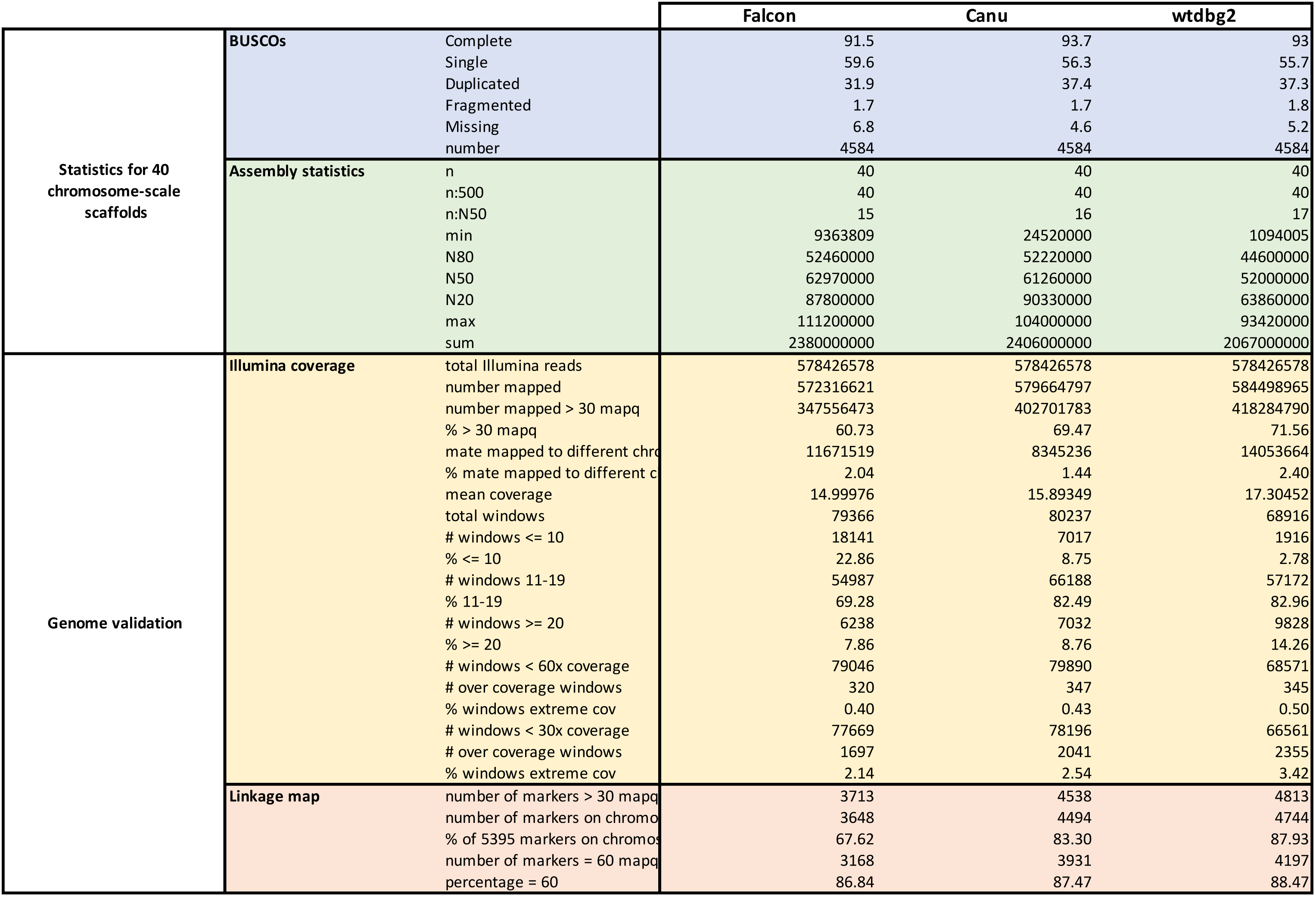
Assembly statistics for the 40 chromosome-scale scaffolds and statistics from assembly validation using Illumina data and linkage map

**Table S3.** BED file for partial and fully collapsed chromosome-scale scaffolds

| scaffold | start | end | description |
| --- | --- | --- | --- |
| 3 | 0 | 31500000 | Partial collapsed chromosome-scale scaffold |
| 6 | 39000000 | 65391737 | Partial collapsed chromosome-scale scaffold |
| 16 | 41500000 | 54216998 | Partial collapsed chromosome-scale scaffold |
| 36 | 32500000 | 43663377 | Partial collapsed chromosome-scale scaffold |
| 21 | 0 | 56862223 | Full collapsed chromosome-scale scaffold |
| 27 | 0 | 46671285 | Full collapsed chromosome-scale scaffold |
| 31 | 0 | 44616205 | Full collapsed chromosome-scale scaffold |
| 34 | 0 | 42609905 | Full collapsed chromosome-scale scaffold |
| 37 | 0 | 36774138 | Full collapsed chromosome-scale scaffold |

